# Mistranslating tRNA variants have anticodon- and sex-specific impacts on *Drosophila melanogaster*

**DOI:** 10.1101/2024.06.11.598535

**Authors:** Joshua R. Isaacson, Matthew D. Berg, Jessica Jagiello, William Yeung, Brendan Charles, Judit Villén, Christopher J. Brandl, Amanda J. Moehring

## Abstract

Transfer RNAs (tRNAs) are vital in determining the specificity of translation. Mutations in tRNA genes can result in the misincorporation of amino acids into nascent polypeptides in a process known as mistranslation. Since mistranslation has different impacts, depending on the type of amino acid substitution, our goal here was to compare the impact of different mistranslating tRNA^Ser^ variants on fly development, lifespan, and behaviour. We established two mistranslating fly lines, one with a tRNA^Ser^ variant that misincorporates serine at valine codons (V➔S) and the other that misincorporates serine at threonine codons (TàS). While both mistranslating tRNAs increased development time and developmental lethality, the severity of the impacts differed depending on amino acid substitution and sex. The V➔S variant extended embryonic, larval, and pupal development whereas the T➔S only extended larval and pupal development. Females, but not males, containing either mistranslating tRNA presented with significantly more anatomical deformities than controls. Mistranslating females also experienced extended lifespan whereas mistranslating male lifespan was unaffected. In addition, mistranslating flies from both sexes showed improved locomotion as they aged, suggesting delayed neurodegeneration. Therefore, although mistranslation causes detrimental effects, we demonstrate that mistranslation also has positive effects on complex traits such as lifespan and locomotion. This has important implications for human health given the prevalence of tRNA variants in humans.

**PLAIN LANGUAGE SUMMARY:** Mutant tRNA genes can cause mistranslation, the misincorporation of amino acids into proteins, and are associated with several human diseases. This study investigated the role of two tRNA variants that cause threonine-to-serine (T➔S) or valine-to-serine (V➔S) substitution. Both variants caused developmental delays and lethality in both sexes and increase prevalence of deformities in females. Surprisingly, female T➔S and V➔S flies experienced increased lifespan and mistranslating males and females showed improved locomotion. These results suggest that mistranslation has both positive and negative effects that depend on the tRNA variant and sex of the fly.

## INTRODUCTION

The translation of nucleotide sequence into protein is a fundamental cellular process that requires a high degree of accuracy. By delivering the correct amino acid to the nascent peptide chain at the ribosome, aminoacylated transfer RNAs (tRNAs) have a principal role in translation fidelity. Aminoacyl-tRNA synthetases (aaRSs) aminoacylate their tRNA substrates with their cognate amino acid (reviewed in Pang *et al*. 2014). Specific bases and motifs within tRNAs, known as identity elements, are recognized by aaRSs to ensure accurate aminoacylation (Hou and Schimmel 1988; Normanly *et al*. 1992; Xue *et al*. 1993). The anticodon, spanning bases 34– 36 and which base-pairs with the mRNA codon, is an identity element for many aaRSs, providing a direct link between the aaRS and codon assignment (Schulman and Pelka 1989; Ruff *et al*. 1991; Jahn *et al*. 1991; Tamura *et al*. 1992; Kholod *et al*. 1997; Giegé *et al*. 1998; Zamudio and José 2018; Giegé and Eriani 2023). If tRNA mischarging does occur, some aaRSs also contain editing domains that deacylate the tRNA (Dock-Bregeon *et al*. 2000; Perona and Gruic-Sovulj 2014; Kuzmishin Nagy *et al*. 2020).

The anticodon is not an identity element for eukaryotic tRNA^Ser^, tRNA^Leu^, and tRNA^Ala^ (McClain and Foss 1988; Hou and Schimmel 1988; Asahara *et al*. 1993; Achsel and Gross 1993; Breitschopf *et al*. 1995; Himeno *et al*. 1997; reviewed in Giegé and Eriani 2023). For tRNA^Ser^, the extended variable arm is the principal identity element (Normanly *et al*. 1992; Achsel and Gross 1993; Lenhard *et al*. 1999). Because only the extended variable arm is required for aminoacylation by SerRS, tRNA^Ser^ variants that contain non-serine anticodons will be serylated and misincorporate serine in place of the anticodon-designated amino acid (Garza *et al*. 1990; Reverendo *et al*. 2014; Berg *et al*. 2017, 2019b; Lant *et al*. 2018; Zimmerman *et al*. 2018; Isaacson *et al*. 2022). Since the ribosome has a limited ability to screen for misacylated tRNAs (Dale *et al*. 2009), tRNA^Ser^ anticodon variants increase mistranslation levels.

Mistranslation has diverse effects on an organism. Mutant aaRSs that cause mistranslation reduce lifespan, impair locomotion, and cause neurodegeneration in *Drosophila melanogaster* (Lu *et al*. 2014), and promote cardiac abnormalities, neurodegeneration and tumor growth in mice (Lee *et al*. 2006; Liu *et al*. 2014; Santos *et al*. 2018). Mistranslating tRNA variants cause developmental deformities in zebrafish and flies (Reverendo *et al*. 2014; Isaacson *et al*. 2022). In human cells, mistranslating tRNAs reduce translation rate and impair clearing of polyQ protein aggregates (Lant *et al*. 2021; Davey-Young *et al*. 2024). Interestingly, mistranslation can also have positive effects (Ribas de Pouplana *et al*. 2014). For example, mistranslation acts as a stress-response mechanism in bacterial, yeast, and human cells to withstand oxidative stress (Santos *et al*. 1999; Netzer *et al*. 2009; Fan *et al*. 2015; Evans *et al*. 2019; Samhita *et al*. 2020).

Despite the effects of mistranslation on cell biology and a recent sequencing study estimating that humans contain ∼66 cytoplasmic tRNA variants per individual including some variants that mistranslate (Berg *et al*. 2019a), the effects of mistranslating tRNAs on multicellular organisms are poorly understood. To address this, we previously created a cytoplasmic tRNA mistranslation model in the fruit fly *Drosophila melanogaster* that misincorporates serine at proline codons (Isaacson *et al*. 2022). Flies containing the mistranslating tRNA variant had increased development time and developmental lethality, more anatomical deformities, and worse climbing performance than flies containing a wild-type serine tRNA. Mistranslating females also presented with more deformities and faster climbing performance decline than males. Since previous work in yeast demonstrated that the effects of mistranslation vary with type of amino acid substitution (Berg *et al*. 2021b; Cozma *et al*. 2023; Davey-Young *et al*. 2024), it is important to determine how different mistranslating tRNA^Ser^ variants impact multicellular physiology. To test this, here we generated tRNA^Ser^ variants that substitute serine at either valine (V➔S) or threonine codons (T➔S) and compared how different types of mistranslation affect flies. Both substitutions extended development time, reduced survival through development, and significantly increased the prevalence of deformities in females. Females from both tRNA^Ser^ lines experienced an increase in lifespan whereas male lifespan was unaffected. Male and female flies containing the variant tRNA^Ser^ genes had improved climbing performance. Thus, mistranslating tRNA variants exert strong positive and negative effects on fruit flies that differ by sex and properties of the mistranslating tRNA variant.

## MATERIALS AND METHODS

### Fly husbandry and stocks

All fly stocks were obtained from the Bloomington *Drosophila* Stock Center and maintained on standard Bloomington recipe food medium (BDSC; Bloomington, Indiana) under a 14:10 light:dark cycle at 24°C and 70% relative humidity.

### Plasmid construction

The shuttle vector used to integrate tRNAs into the *D. melanogaster* genome is pattB, which was a kind gift from Bischof *et al*. (2012, DGRC #1420). The *Not*I site within pattB was removed through digestion and blunting with the Klenow fragment of DNA polymerase, creating pattB-*Not*IΔ. A tRNA^Ser^_UGG, G26A_ gene (a variant of FlyBase ID: FBgn0050201, Öztürk-Çolak *et al*. 2024), along with ∼300 bp of upstream and downstream sequence, was flanked with FRT sites and synthesized by Integrated DNA Technologies, Inc. The tRNA sequence within the FRT sites was bookended by *Not*I sites, enabling swapping the tRNA by cloning in a new tRNA gene as a *Not*I fragment, and the entire FRT-tRNA-FRT fragment was flanked by *Eco*RI and *Bam*HI sites. The synthesized fragment was cloned into pattB-*Not*IΔ as an *Eco*RI/*Bam*HI fragment, creating pattB-*Not*IΔ/pUCIDT (Figure S1).

The serine tRNA variant containing a valine AAC anticodon and G26A mutation (tRNA_AAC_^Ser^) was made through two-step PCR using tRNA_UGA_^Ser^ (FlyBase ID: FBgn0050201) from genomic DNA as a template. The primers tSerAAC_F/tSerDS and tSerAAC_R/tSerUS were used in the first round and products from the first round were amplified using outside primers tSerUS/tSerDS during the second round (all primer sequences are listed in Table S1). Second round PCR products were cloned into pGEM^®^-T Easy (Promega) and sequenced. Correct plasmids were digested with *Not*I and the tRNA fragment cloned into pattB-*Not*IΔ/pUCIDT to flank the tRNA with FRT sites. An identical procedure was used to create the serine tRNA variant containing a threonine AGT anticodon and G26A mutation (tRNA_AGU_^Ser^) using primers tSerAGU_F/tSerDS and tSerAGU_R/tSerUS in the first round.

### Creating mistranslating stocks

Mistranslating tRNAs were integrated into flies by injecting plasmids into *D. melanogaster* embryos from BDSC stock # 24872 (*y^1^ M{RFP[3xP3.PB] GFP[E.3xP3]=vas-int.Dm}ZH-2A w*; PBac{y^+^-attP-3B}VK00037*), which expresses phiC31 (ΦC31) in the germ line and contains an *attP40* site in the left arm of the second chromosome. The *attP40* landing site was selected as it is relatively inert while allowing for strong expression of transgenes (Markstein *et al*. 2008). The injection protocol has been described (Isaacson 2018). Transgenic flies were identified through their mini-white eye colour and balanced using BDSC stock # 3703 (*w^1118^/Dp(1;Y)y^+^; CyO/nub^1^ b^1^ sna Sco lt^1^ stw^3^; MKRS/TM6B, Tb^1^*) to create stocks of the genotype *w^1118^*; *P{CaryP}-attP40[w^mw+^=pattB-tRNA]*/*CyO*; *MKRS*/*TM6B*. DNA was extracted from parents of the final cross, PCR amplified using the primer set pattB-tRNA-Ver_F/pattB-tRNA-Ver_R, and sequenced to confirm accuracy of the inserted tRNA.

### Creating FLP-out controls

Flanking the inserted tRNA with FRT sites oriented in the same direction allowed removal of the inserted tRNA in the presence of flippase (Gronostajski and Sadowski 1985). To remove the tRNA and create control lines, flies containing tRNA_AAC_^Ser^ or tRNA_AGU_^Ser^ were crossed to a UAS-FLP line (BDSC stock # 4540: *w*; P{w^+mC^=UAS-FLP.D}JD2*) and a germ-line specific *nanos*-Gal4 line (BDSC stock # 4937: *w^1118^; P{w^+mC^=GAL4::VP16-nanos.UTR}CG6325^MVD1^*). Offspring were crossed to each other and removal of the tRNA in both parents was confirmed by PCR and sequencing using primer set FRT-tRNA-Ver_F/FRT-tRNA-Ver_R. Successful tRNA FLP-out lines were then crossed back to stock #3703 to create control lines of the following genotype: *w^1118^*; *P{CaryP}-attP40[w^mw+^=pattB-FLP-out]*/*CyO*; *MKRS*/*TM6B*. Control lines for tRNA_AAC_^Ser^ are referred to as tRNA_AAC_^Ser^ -FLP and control lines for tRNA^Ser^_AGU_ are referred to as tRNA_AGU_^Ser^-FLP.

### Mass spectrometry

Five replicates of twenty pupae or ten adult flies were collected from each genotype and lysed in 8 M urea, 50 mM Tris, 75 mM NaCl, pH 8.2 by beating with 0.5 mm glass beads at 4°C and protein concentration was determined by bicinchoninic acid assay (Pierce, ThermoFisher Scientific). Protein was reduced with 5 mM dithiothreitol for 30 minutes at 55°C, alkylated with 15 mM iodoacetamine for 30 minutes at room temperature in the dark and the alkylation was quenched with an additional 5 mM dithiothreitol for 30 minutes at room temperature. For each sample, 50 µg of protein was diluted four-fold with 50 mM Tris pH 8.9 and digested for 4 hours at 37°C with 1.0 µg endoproteinase Lys-C (Wako Chemicals). Digestions were acidified to pH 2 with trifluoroacetic acid and peptides were desalted by solid phase extraction over Empore C18 stage tips (Rappsilber *et al*. 2007) and dried by vacuum centrifugation.

Peptides were resuspended in 4% acetonitrile, 3% formic acid and subjected to liquid chromatography coupled to tandem mass spectrometry (LC-MS/MS) on a tribrid quadrupole Orbitrap mass spectrometry (Orbitrap Eclipse; ThermoFisher Scientific) operated in data dependent acquisition mode as described in Cozma *et al*. (2023).

MS/MS spectra were searched against the *D. melanogaster* protein sequence database (version r6.09; downloaded from FlyBase in 2016, Öztürk-Çolak *et al*. 2024) using Comet (release 2015.01; Eng *et al*. 2013). The precursor mass tolerance was set to 20 ppm. Constant modification of cysteine carbamidomethylation (57.0215 Da) and variable modification of methionine oxidation (15.9949 Da) and protein N-terminal lysine acetylation (42.0102 Da) were used for all searches. A variable modification of valine to serine (-12.0364 Da) or threonine to serine (-14.0156 Da) were used for the respective mistranslating tRNA and control samples. A maximum of two of each variable modification were allowed per peptide. Search results were filtered to a 1% false discovery rate at the peptide spectrum match (PSM) level using Percolator (Käll *et al*. 2007). The mistranslation frequency was calculated using the unique mistranslated peptides for which the non-mistranslated sibling peptide was also observed. The frequency is defined as the counts of mistranslated peptides, where serine was inserted for valine or threonine, divided by the counts of all peptides containing valine or threonine, respectively, and expressed as a percentage. At the codon level, the decoding specificity of each tRNA^Ser^ variant was determined using a custom Perl script described in Cozma *et al*. (2023). Briefly, codons were mapped back to wild-type and mistranslated residues for all peptides with only one possible substitution event to allow for accurate localization of the mistranslated residue. Mistranslation frequency at each codon was determined from counts as above.

### Development assay

Approximately 250 flies from each of the four genotypes were placed into fly cages and allowed to lay eggs for one hour. Equal numbers of eggs were collected from each plate and checked every 12 hours to record progress through each of the following developmental stages: egg hatching into larva, larva pupating into pupa, and adult eclosing from pupa. In total, 200 eggs from each genotype were collected. Sex and zygosity of adults were recorded.

### Scoring for deformities

Virgin, heterozygous flies from the two mistranslating lines and their corresponding controls were collected within ∼8 hours of eclosion and scored for deformities in adult legs (limbs gnarled or missing segments), wings (blistered, absent, fluid-filled, or abnormal size), or abdomen (fused or incomplete tergites). Flies collected before wing expansion were excluded. Sex and type of deformity was recorded. Flies that had multiple deformities had each recorded. For the threonine lines, 591 tRNA^Ser^_AGU_ (287 males and 304 females) and 550 tRNA^Ser^_AGU_-FLP (276 males and 274 females) flies were scored. For the valine lines, 723 tRNA^Ser^_AAC_ (373 males and 350 females) and 552 tRNA^Ser^_AAC_-FLP (282 males and 270 females) flies were scored. Deformities were photographed through the lens of a stereomicroscope using a Samsung Galaxy S8 camera.

### Longevity assays

Equal numbers of adult, virgin flies of each sex were collected from all lines within 8 hours of eclosion and placed in new food vials (101 flies for each threonine line and 119 flies for each valine line). Flies with deformities were noted but still used in the assay. Flies were transferred to new food every three days and deaths were recorded. If dead flies were found in a vial known to contain a fly with a deformity, the dead fly was examined for deformities.

### Climbing assays

Climbing assays were conducted on the flies in the longevity assay. The day before testing, flies were transferred to fresh food. The number of flies that reached a goal line 5 cm above the surface of the food within 10 seconds were recorded. Each vial was tested three times. Climbing performance calculated as the percentage of successful flies out of the total number of flies in the vial. Flies were tested 30, 51, and 72 days after eclosion. Only nondeformed flies were considered when recording the total number of flies (e.g. a vial with six flies but one deformed fly was treated as containing five flies).

### Statistical analyses

Statistical analyses were performed using R Studio v1.2.5001. Analyses used for comparisons were: *t*-test (frequency of T➔S misincorporation between tRNA_AGU_^Ser^ and tRNA^Ser^_AGU_-FLP, or V➔S misincorporation between tRNA^Ser^_AAC_ and tRNA_AAC_^Ser^-FLP); Wilcoxon rank-sum tests (developmental time data, corrected using Holm-Bonferroni’s method); and Fisher’s exact tests (survival between developmental stages, proportion of deformities, and climbing performance, all corrected using Holm-Bonferroni’s method). Fly longevity was quantified and compared using the “survminer” R package (Kosinski *et al*. 2020) and log-rank tests corrected using Holm-Bonferroni’s method. Because the climbing assays were performed on flies undergoing the longevity assay, climbing assay and longevity assay *P*-values were corrected together. All raw data can be found in Supplemental file S1.

### Data and Reagent Availability

Fly lines and plasmids are available upon request. The authors affirm that all data necessary for confirming the conclusions of the article are present within the article, figures, and supplemental material. Supplemental File S1 contains all raw data and descriptions of the mass spectrometry files. Supplemental File S2 contains supplemental methods, figures, and tables. Supplemental File S3 contains R code used to analyze all raw data. The mass spectrometry proteomics data have been deposited to the ProteomeXchange Consortium via the PRIDE partner repository (Perez-Riverol *et al*. 2019) with the dataset identifier PXD052492.

## RESULTS

### Creating mistranslating fly lines

To assess the impacts of different mistranslating tRNA variants on *Drosophila melanogaster*, we mutated the anticodon of the gene encoding *D. melanogaster* tRNA^Ser^ (FlyBase ID: FBgn0050201) to AAC or AGT to engineer tRNA^Ser^ variants that misincorporate serine at valine (V➔S) or serine at threonine codons (T➔S), respectively. The tRNA sequence also included a G26A base change to remove a key modification site in tRNA^Ser^ (Boccaletto *et al*. 2022). The G26A change causes increased degradation of the tRNA^Ser^ variants through the rapid tRNA decay pathway and ensures mistranslation occurs at tolerable levels based on work in yeast (Dewe *et al*. 2012; Berg *et al*. 2021a). We flanked the tRNA variant with FRT sites to allow flippase driven in the germ line to excise the tRNA from germ cells and produce control offspring with no copies of the inserted tRNA. The control lines (tRNA_AAC_^Ser^-FLP and tRNA^Ser^_AGU_-FLP) share a genetic background with their corresponding mistranslating line (tRNA_AAC_^Ser^ and tRNA_AGU_^Ser^). The presence of the tRNA variants in the experimental lines and their absence in the controls was confirmed through PCR and sequencing.

To determine frequency of T➔S or V➔S mistranslation, we analyzed the proteome of pupae and adults from mistranslating *Drosophila* lines and their respective control using mass spectrometry. We define mistranslation frequency as the number of unique peptides observed where valine or threonine is replaced with serine relative to the total number of peptides observed with valine or threonine (see Methods for details). Levels of translation are relatively high in pupae and this developmental stage determines adult neuronal and skeletomuscular structures (Mitchell *et al*. 1977; Mitchell and Petersen 1981; Truman and Bate 1988; Truman 1990). In pupae, both lines had significantly higher mistranslation frequencies than control lines, though the frequency of T➔S mistranslation was higher than V➔S (Figure 1A, B). As the frequency of V➔S mistranslation was low, we manually examined peptide spectra to confirm mistranslation. Spectra for mistranslated and wild-type peptides are shown in Figure S2.

**Figure 1.**
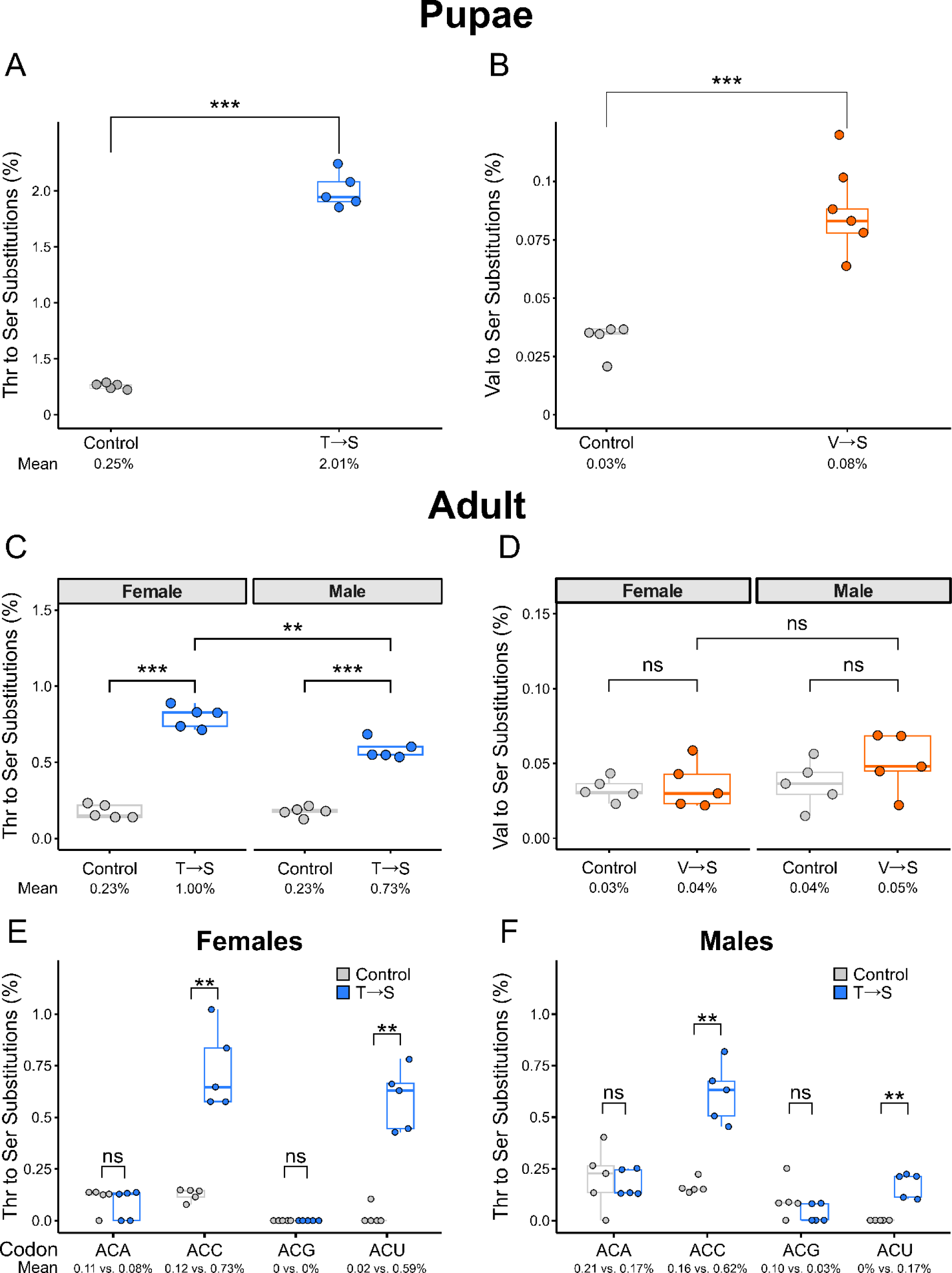
Mistranslation frequency of pupae and adult flies containing tRNA^Ser^_AGU_ (T➔S) or tRNA^Ser^_AAC_ (V➔S) as determined from whole-proteome mass spectrometry. **A)** Frequency of T➔S mistranslation in tRNA_AGU_^Ser^-FLP (control) and tRNA_AGU_^Ser^ (T➔S) pupae (n = 5 replicates of 20 pupae each). Numbers below the x-axis represent average mistranslation frequency. **B)** Frequency of V➔S mistranslation in tRNA_AAC_^Ser^-FLP (control) and tRNA_AAC_^Ser^ (V➔S) pupae. **C)** Frequency of T➔S mistranslation in 1–3-day old female or male adult flies containing tRNA^Ser^_AGU_-FLP (control) or tRNA_AGU_^Ser^ (T➔S) (n = 5 replicates of 10 flies each). **D)** Frequency of V➔S mistranslation in female or male adult flies containing tRNA_AAC_^Ser^-FLP (control) or tRNA_AAC_^Ser^ (V➔S). Note the difference in y-axis scale between panels. **E)** Frequency of T➔S mistranslation at all four ACN codons in adult female tRNA_AGU_^Ser^ (T➔S) compared to adult female tRNA^Ser^_AGU_-FLP (control) flies (n = 5 replicates of 10 flies each). Numbers below the x-axis represent average misincorporation frequency. **F)** Same as **E)** but for male adult tRNA_AGU_^Ser^ flies. Genotypes were compared with a *t*-test. “ns” *P* > 0.05, “**” *P* < 0.01; “***” *P* < 0.001.

Mistranslation frequencies decreased in adulthood for both T➔S and V➔S lines compared to pupation (Figure 1C, D). The frequency of T➔S mistranslation was significantly higher for both female and male adults compared to controls (Figure 1C). Interestingly, T➔S females mistranslated significantly more often than T➔S males. Observed frequencies of V➔S mistranslation for adult females and males were not significantly higher than control flies (Figure 1D). However, given the low frequency of mistranslation observed in V➔S pupae we infer that mistranslation is occurring in V➔S adults but at a frequency below the threshold of detection. These results show that our new fly lines experience mistranslation, and that mistranslation frequencies vary by developmental stage and sex.

We also determined which codons were being mistranslated in T➔S female and male flies (Figure 1E, F). The fully complementary ACU codon was significantly mistranslated, whereas the 3’ mismatched ACG was not. Both ACA and ACC codons are expected to be decoded through wobble base pairing if A34 of tRNA_AGU_^Ser^ is deaminated to inosine (Crick 1966; Agris *et al*. 2018; Boccaletto *et al*. 2022). Interestingly, mistranslation was significantly higher at ACC codons, but not ACA, in T➔S flies compared to controls. We note that *D. melanogaster* contain native tRNA^Thr^_CGU_ and tRNA^Thr^_UGU_ to decode ACG and ACA codons.

### Mistranslation extends developmental time and causes lethality

To determine how T➔S and V➔S mistranslation affects development, we collected 200 eggs from each of the mistranslating lines and their controls and counted the number of individuals that survived to larval, pupal, and adult stages. We also measured the time until eggs hatch into larvae, larvae pupate into pupae, and pupae eclose into adults to determine if developmental delays occurred. Significantly fewer T➔S individuals hatched and eclosed compared to controls (Figure 2A), whereas survival of V➔S flies was only significantly reduced during the adult life stage transition (Figure 2B). Measurement of development time revealed that T➔S flies took significantly longer to pupate and eclose compared to controls (Figure 3B, C). V➔S individuals took significantly longer to reach all life stage transitions compared to control flies (Figure 3D-F). Overall, both tRNA^Ser^ variants caused developmental delays and lethality, though different mistranslating tRNA variants affected different life stages.

**Figure 2.**
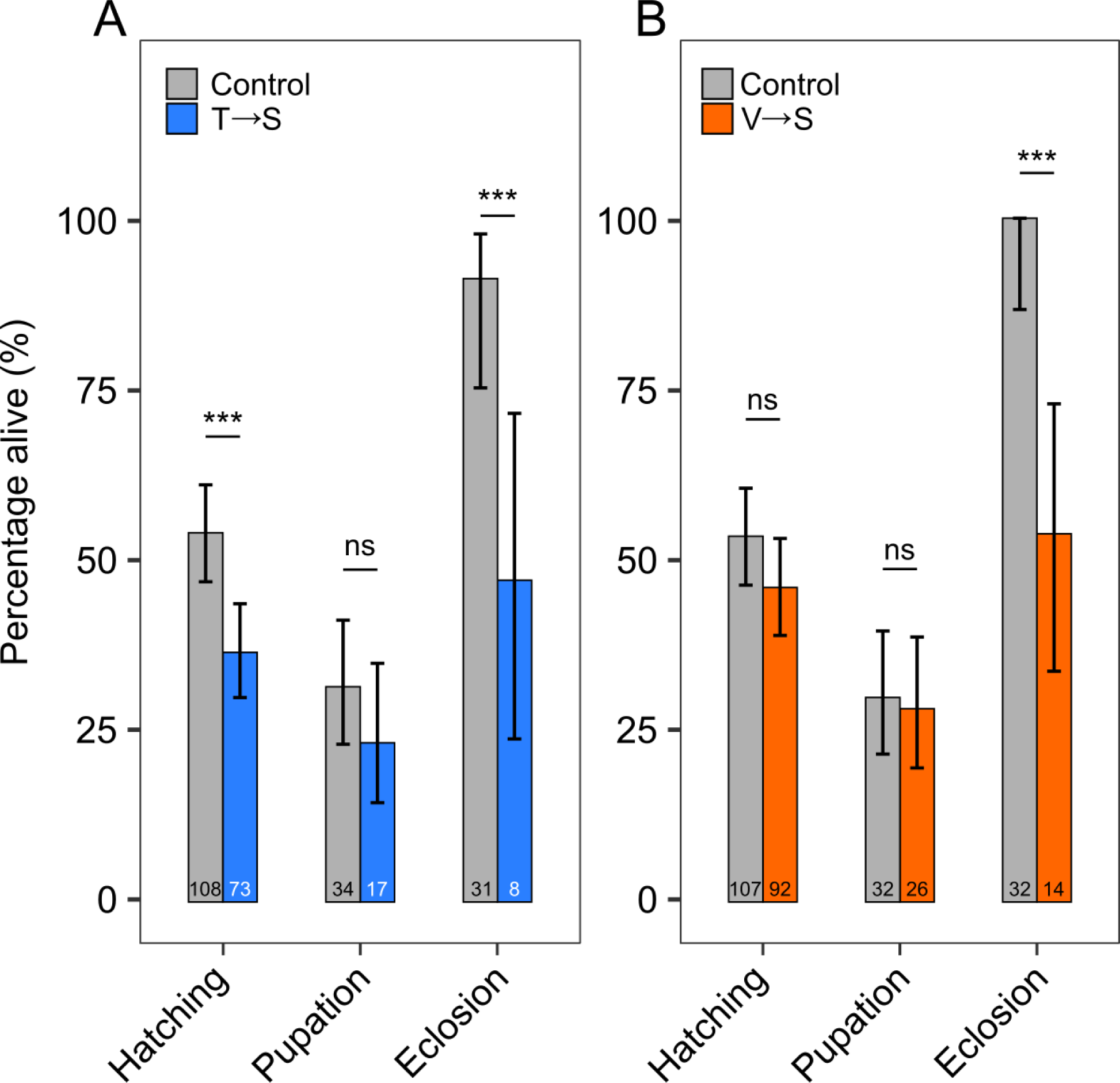
Flies containing tRNA^Ser^_AGU_ (T➔S) and tRNA^Ser^_AAC_ (V➔S) experience developmental lethality. Bars show the percentage of 200 embryos from **A)** tRNA^Ser^_AGU_ (T➔S) and tRNA^Ser^_AGU_-FLP (control) or **B)** tRNA_AAC_^Ser^ (V➔S) and tRNA_AAC_^Ser^-FLP (control) individuals that successfully hatched, pupated, and eclosed. Numbers within the bars indicate the number of embryos that survived beyond that life stage transition and percentages describe the number of survivors from the previous stage that survived beyond the current transition. Error bars represent the 95% confidence interval of the proportion. Survival rates were compared using Fisher’s exact test corrected using Holm-Bonferroni’s method. “ns” *P* ≥ 0.05; “***” *P* < 0.001.

**Figure 3.**
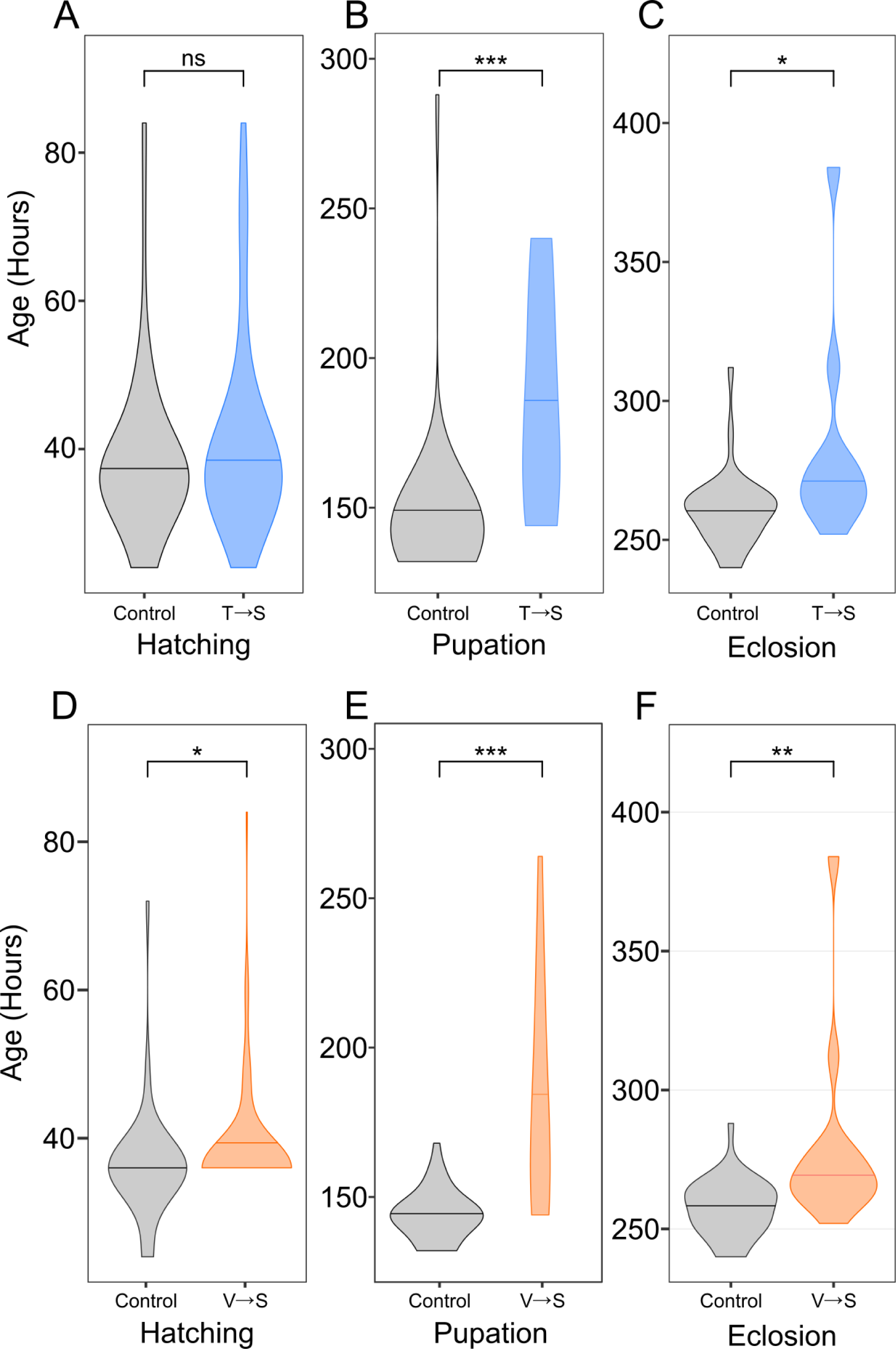
tRNA^Ser^_AGU_ (T➔S) and tRNA^Ser^_AAC_ (V➔S) extended fly development. **A)** Violin plot representing the distribution of total time it took tRNA_AGU_^Ser^ (T➔S) or tRNA_AGU_^Ser^-FLP (control) one-hour old embryos to hatch into larvae, **B)** pupate into pupae, and **C)** eclose into adult flies. **D)** Violin plot representing total time it took tRNA_AAC_^Ser^ (V➔S) or tRNA_AAC_^Ser^-FLP (control) one-hour old embryos to hatch into larvae, **E)** pupate into pupae, and **F)** eclose into adult flies. Horizontal bars within the plot represent the median of the distribution. Sample size is identical to the values within the corresponding bars in Figure 2. Statistical comparisons were performed using Wilcoxon rank-sum tests corrected using Holm-Bonferroni’s method. “ns” *P* ≥ 0.05; “*” *P* < 0.05; “**” *P* < 0.01; “***” *P* < 0.001.

Next, we determined if there was a difference in sex or zygosity distribution among flies that reached the adult stage. Because there were only 23 surviving adults available to score from the tRNA^Ser^ variant lines (one adult was lost during transfer), data from the two mistranslating lines were pooled to assess if mistranslation caused any general trends. Sex distribution of surviving adults was roughly 50% for both tRNA^Ser^ variant lines and their controls (Table 1). Interestingly, 91% of adult flies containing a tRNA^Ser^ variant were heterozygotic in comparison to 66% for the control flies, the latter matching the 2:1 heterozygote:homozygote ratio expected. This suggests that two copies of the tRNA^Ser^ variant are poorly tolerated by flies and thus few homozygous flies reach the adult stage.

**Table 1.**
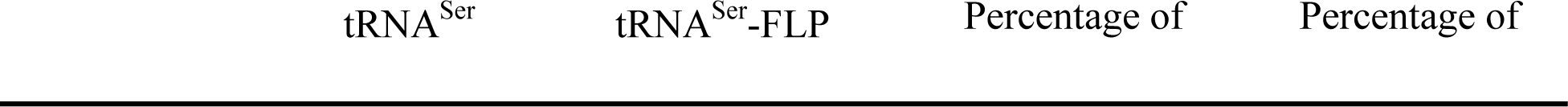
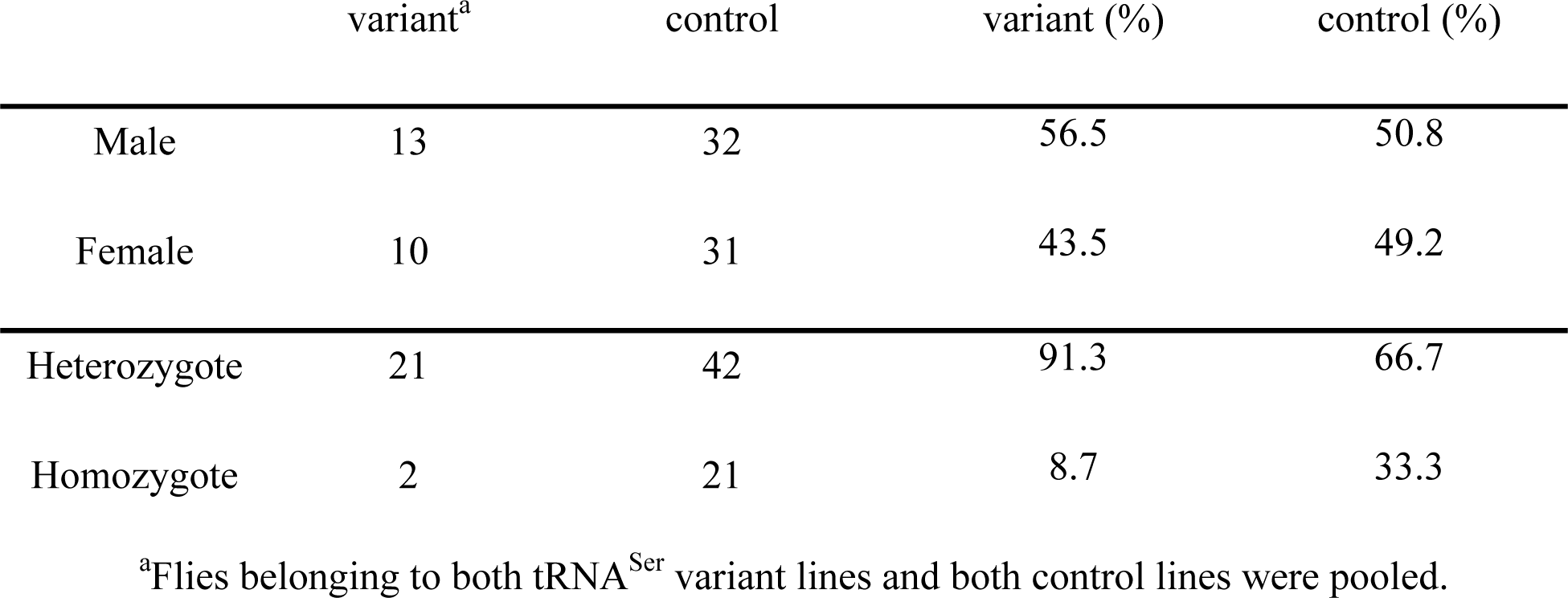
Categorization of adult flies that survived the developmental assay by sex and zygosity.

### Mistranslation causes deformities in adult female flies

We previously showed that flies containing a tRNA^Ser^ variant that causes P➔S mistranslation increases the prevalence of anatomical deformities with female flies containing this tRNA variant twice as likely to present with at least one deformity compared to males (Isaacson *et al*. 2022). We therefore determined if T➔S or V➔S mistranslating tRNA variants cause deformities in flies and if so, whether it is sex specific. Adult heterozygous flies from all four lines were separated by sex and scored for leg, wing, and tergite defects (Figure 4A-D).

**Figure 4.**
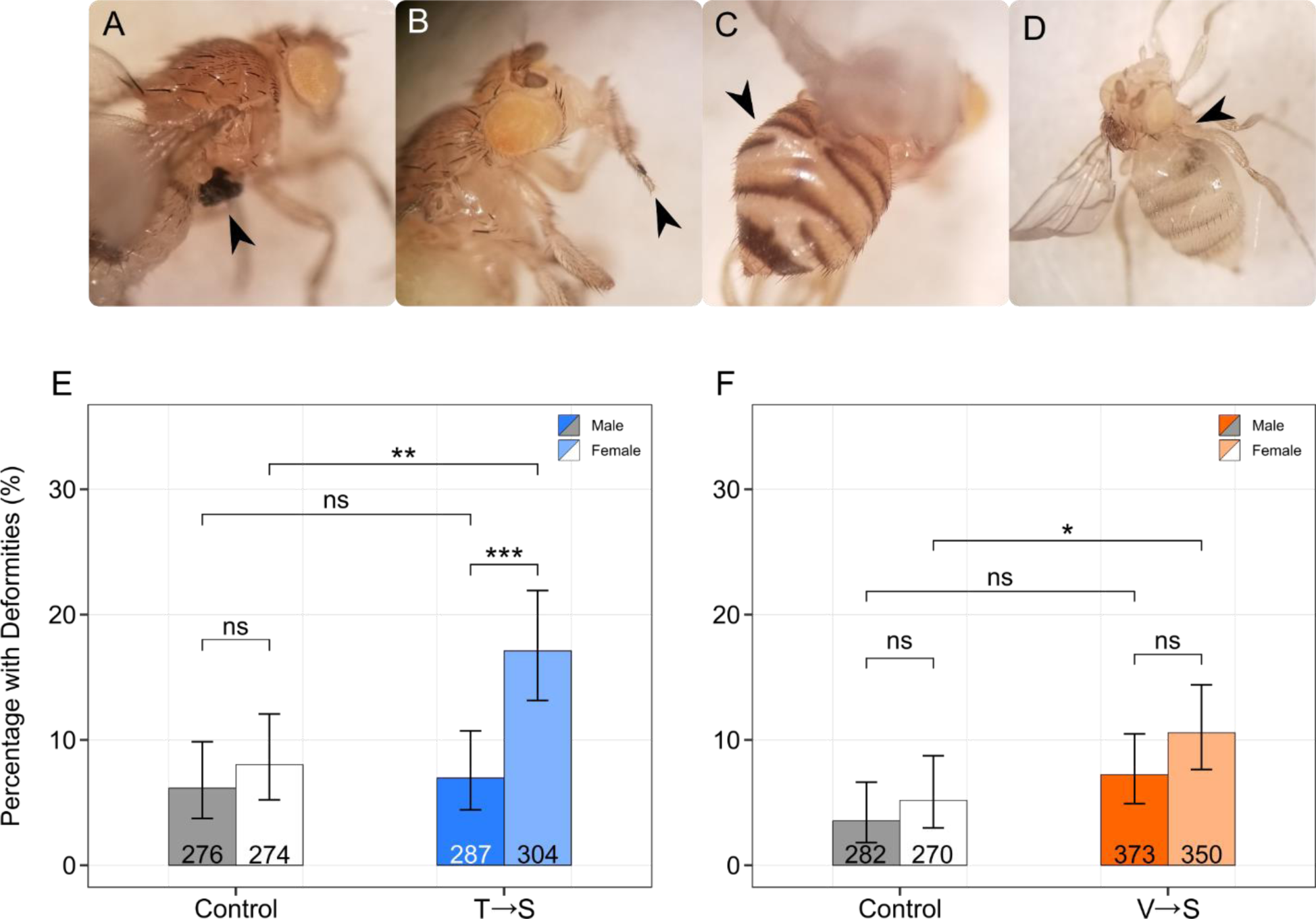
tRNA^Ser^_AGU_ (T➔S) and tRNA^Ser^_AAC_ (V➔S) increase prevalence of deformities in adult female flies. **A)** Example of a fly with a malformed leg, **B)** missing metatarsal, **C)** misfused tergites or **D)** a missing wing. **E)** Percentage of male or female tRNA^Ser^_AGU_ (T➔S) or tRNA^Ser^_AGU_-FLP (control) flies presenting with at least one deformity. Values within bars describe the number of flies examined for deformities. **F)** Percentage of male or female tRNA^Ser^_AAC_ (V➔S) or tRNA_AAC_^Ser^-FLP (control) flies presenting with at least one deformity. Groups were compared using Fisher’s exact test and corrected for multiple comparisons using Holm-Bonferroni’s method. Error bars represent the 95% confidence interval of the proportion. “ns” *P* ≥ 0.05; “*” *P* < 0.05; “**” *P* < 0.01; “***” *P* < 0.001.

Female T➔S flies presented with deformities more than twice as often as control females and mistranslating males (Figure 4E). In contrast, male T➔S flies eclosed with a similar number of deformities as male control flies. V➔S females also presented with significantly more deformities than control females (Figure 4F). An increased fraction of male V➔S flies presented with deformities compared to male control flies, but this difference was not statistically significant. Likewise, V➔S females had a greater tendency toward deformities than V➔S mistranslating males but the difference was not statistically significant. These results, in combination with our previous results using P➔S flies (Isaacson *et al*. 2022), indicate that females are more susceptible to the proteotoxic effects of tRNA-induced mistranslation during development.

### Mistranslating tRNA variants increase female fly lifespan

Given the dramatic effects of mistranslation on development time and survival, we next tested whether mistranslation affects the lifespan of adult flies. Equal numbers of heterozygous virgin males and females from each mistranslating tRNA^Ser^ variant line and its control were collected and transferred to new food vials every three days. Dead flies were recorded and removed during transfer, and survival curves were calculated. In total, 101 male and female flies were analyzed for each of the tRNA_AGU_^Ser^ (T➔S) and tRNA_AGU_^Ser^-FLP control lines and 119 were analyzed for the tRNA_AAC_^Ser^ (V➔S) and tRNA_AAC_^Ser^-FLP control lines. As expected, virgin female lifespan was longer than male lifespan for both T➔S and V➔S flies (Figure 5A, B), as virgin females tend to live longer than virgin males (Ziehm *et al*. 2013). Neither T➔S nor V➔S males flies experienced a change in lifespan compared to control male flies (Figure 5C, D). Interestingly, both T➔S and V➔S females lived longer than control females (Figure 5E, F). These results persist when deformed flies are removed from analysis (Supplemental File S3). Our results demonstrate that two different mistranslating tRNA^Ser^ variants extend female *D. melanogaster* lifespan without impacting male longevity.

**Figure 5.**
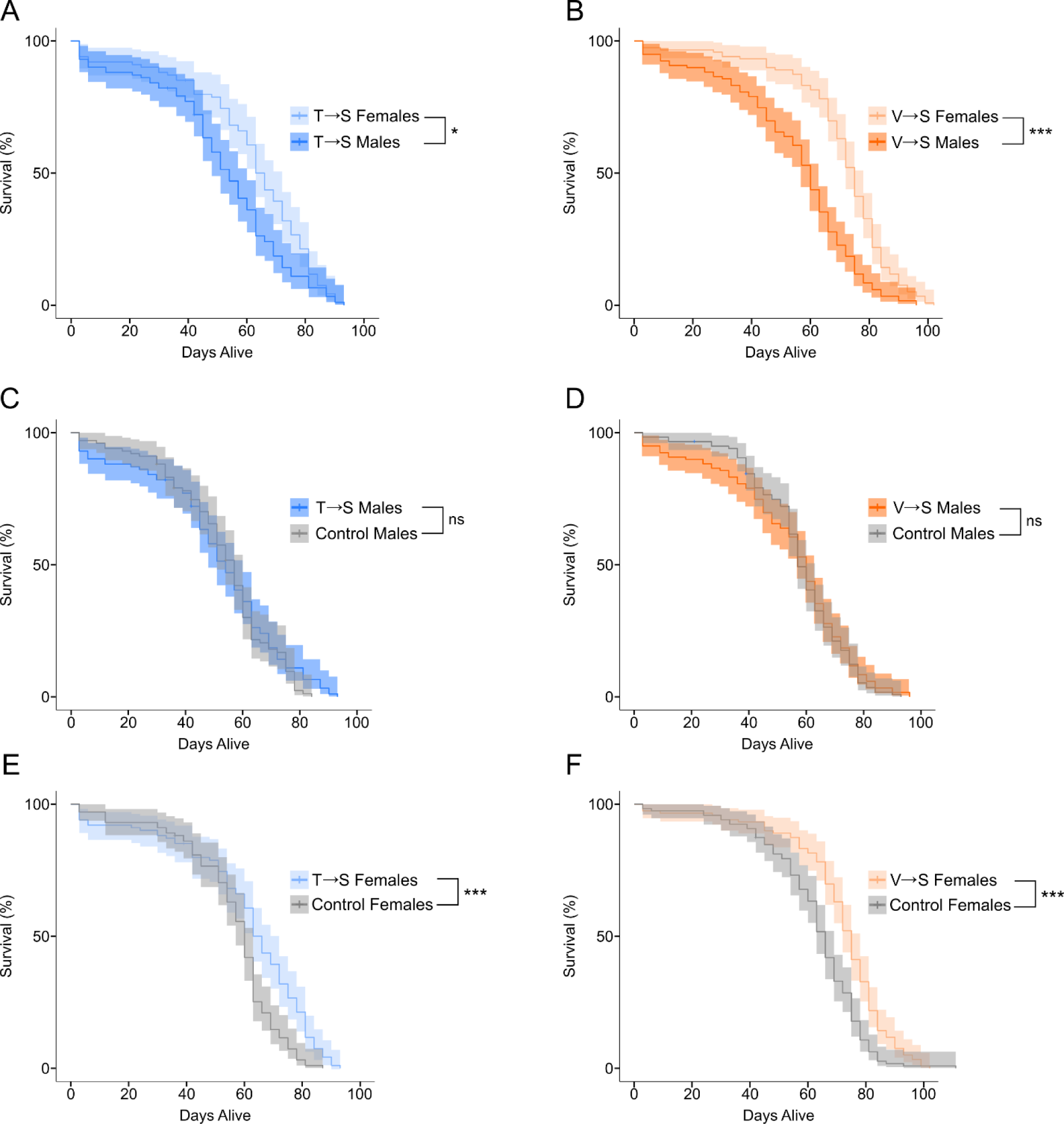
Mistranslating tRNA variants increase female Drosophila melanogaster lifespan. Adult virgin flies were collected within eight hours of eclosion and transferred to new food and scored for survival every three days. 101 male and female tRNA^Ser^_AGU_ (T➔S) and tRNA^Ser^_AGU_-FLP (control) flies were collected, and 119 male and female tRNA^Ser^_AAC_ (V➔S) and tRNA^Ser^_AAC_-FLP (control) flies were collected. Vertical ticks along the line represent censored observations (e.g escaped flies). Kaplan-Meier survival curves are shown with the shaded region representing the 95% confidence interval of survival probability. Survival curves were statistically compared using log-rank tests corrected using Holm-Bonferroni’s method. “ns” *P* ≥ 0.05; “*” *P* < 0.05; “***” *P* < 0.001.

### Mistranslating tRNA^Ser^ variants improve fly climbing performance

Fly performance in negative geotaxis assays, also known as climbing assays, is commonly used as an indicator of neurodegeneration (e.g., Feany and Bender 2000; Song *et al*. 2017; Aggarwal *et al*. 2019). We conducted climbing assays on 30, 51, and 72-day old adult virgin flies that were undergoing the longevity assay (Figure 6). Flies were tested for their ability to climb 5 cm in 10 seconds with each vial tested three times. Surprisingly, both male and female T➔S flies climbed significantly better than their control flies at 30 days of age (Figure 6A). Improvements to climbing performance were observed in V➔S flies as well, but only for females which climbed significantly better than control females and V➔S males (Figure 6B). As an additional method to quantify neurodegeneration, thirty-day old flies were scored for a rough-eye phenotype (Sang and Jackson 2005; Prüßing *et al*. 2013). We observed no difference in the amount of visible neurodegeneration between mistranslating and control flies at this age (Figure S3), which agrees with the climbing assay suggesting that V➔S and T➔S mistranslation does not lead to neurodegeneration in 30-day old flies.

**Figure 6.**
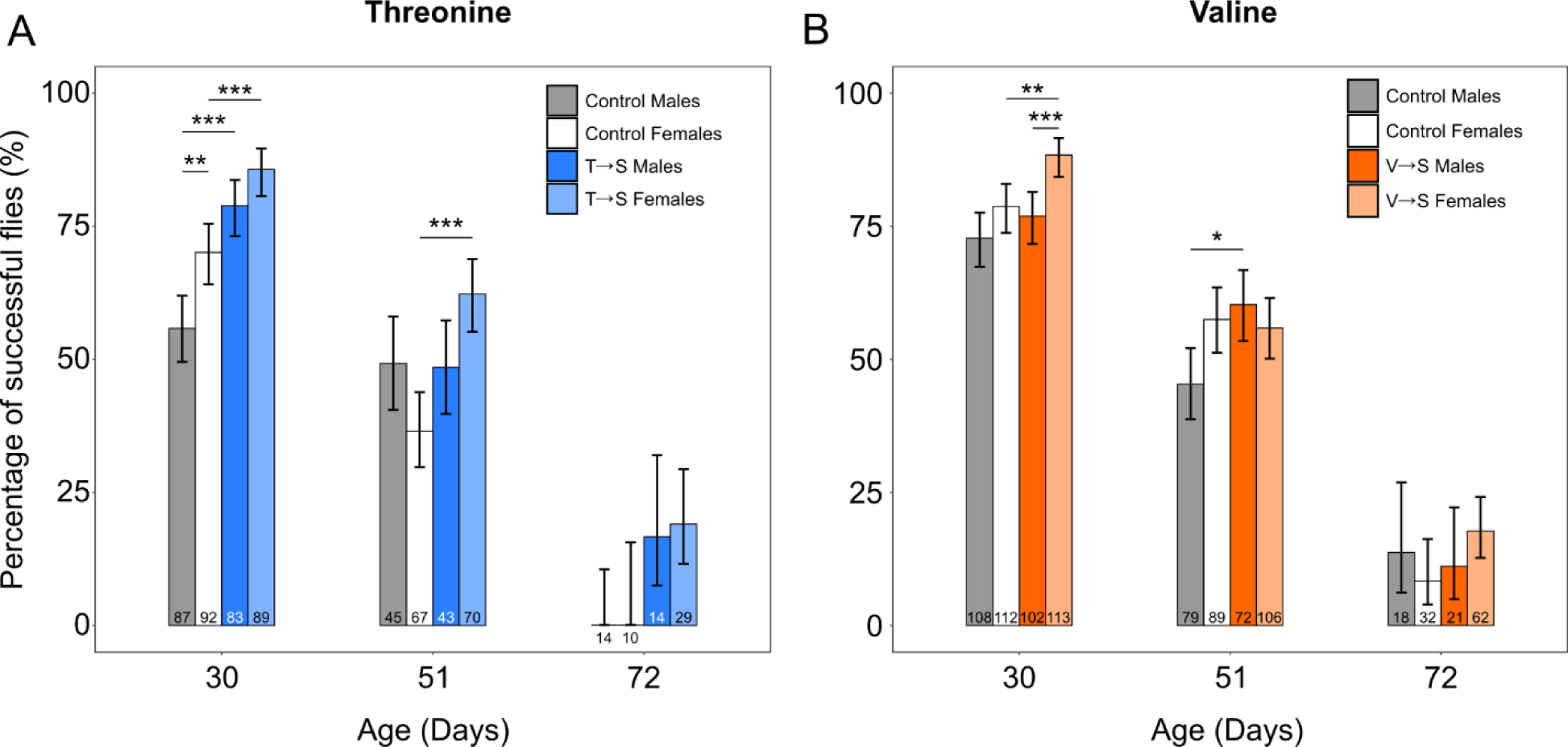
Adult flies containing mistranslating tRNA^Ser^ variants have similar or improved climbing performance compared to control. Each bar represents the percentage of flies from the specified genotype that successfully reached a 5 cm goal line in 10 seconds. All flies were tested three times. The numbers within or below bars represent the number of flies of that genotype and age that were tested. **A)** Climbing performance of tRNA_AGU_^Ser^ (T➔S) or control (tRNA_AGU_^Ser^-FLP) male and female flies at 30, 51, and 72 days of age. **B)** Climbing performance of tRNA^Ser^_AAC_ (V➔S) or control (tRNA^Ser^_AAC_-FLP) male and female flies at 30, 51, and 72 days of age. Performance was compared between groups using Fisher’s exact test and *P*-values were corrected for multiple comparisons using Holm-Bonferroni’s method. Error bars represent the 95% confidence interval of the proportion. Only significant comparisons are shown. “*” *P* < 0.05; “**” *P* < 0.01; “***” *P* < 0.001.

There was substantial die-off in all lines by day 51 which affected statistical power. This decrease in sample size was most pronounced in male T➔S and control flies, as nearly half of the flies alive at day 30 had died by day 51. Despite this reduction in power, 51-day old female T➔S flies still climbed significantly better than control females (Figure 6A), and V➔S males climbed significantly better than control males (Figure 6B). At 72 days, no groups showed significantly different performance, though we note that no control T➔S males or females successfully reached the 5 cm goal line whereas ∼15% of mistranslating T➔S male and females reached the goal (Figure 6A). These findings suggest that T➔S and V➔S mistranslation do not cause neurodegeneration and may instead confer neuroprotective effects at the ages tested.

## DISCUSSION

Mistranslating tRNA^Ser^ variants have complex and sex specific effects on *Drosophila melanogaster*. tRNA variants that mistranslate T➔S and V➔S caused developmental lethality, extended developmental time, and improved adult climbing performance. For female flies, T➔S and V➔S mistranslation increased the prevalence of deformities, however they also extended lifespan.

Based on codon usage and the number of competing tRNAs, we would expect the T➔S variant to cause less mistranslation than the V➔S variant (Supplemental Table S2). However, we instead observed lower mistranslation levels from the V➔S variant compared to the T➔S variant. Both tRNA^Ser^ variants were integrated into the same location on chromosome 2L and maintained as a single copy, so differences in mistranslation frequency are likely not due to position effects or copy number. We note that the levels of mistranslation that we report are based on steady-state protein levels and the limit of detection of the mass spectrometer. For this reason, our estimates of mistranslation frequency are likely an underestimate since some mistranslation events would result in protein turnover. One might predict that the conservative T➔S change would be less deleterious to protein structure, thus minimally impacting protein turnover and increasing the observed mistranslation frequency compared to V➔S mistranslation.

The V➔S and T➔S variants replicated some but not all of our previous results with tRNA^Ser^_UGG_ (P➔S) (Isaacson *et al*. 2022). The prevalence of mistranslation we previously observed for the P➔S variant (∼0.6%) was intermediate to that observed for V➔S and T➔S variants. While all three tRNA variant lines extended development time, increased developmental lethality, and increased deformities in female flies, male and female V➔S and T➔S flies climbed better than their controls while the P➔S flies had impaired climbing performance. This difference could result if proteins involved in neuromuscular function are more sensitive to serine substitution at proline than serine substitution at valine or threonine through mechanisms that could include the prevalence of functionally important prolines in proteins essential for this function.

### Different male and female response to mistranslation

Females experienced stronger positive and negative effects of mistranslation for the three tRNA^Ser^ variants we have tested (this study and Isaacson *et al*. 2022). Females containing the T➔S variant had an observed mistranslation frequency greater than males, which likely influences the sex differences noted in this study. In addition, female susceptibility to mistranslation could be affected by increased developmental requirements, as females are larger, develop faster, and invest more resources into their gametes than male flies (Bonnier 1926; Bakker 1959; Fredriksson *et al*. 2012). Certain environmental conditions also extend lifespan primarily in one sex. For example, dietary restriction, particularly restriction of protein intake, extends lifespan of female flies more than males (Nakagawa *et al*. 2012; Regan *et al*. 2016; Garratt 2020). It would be interesting to test how mated *vs.* virgin flies respond to mistranslation, as mating status heavily impacts fly lifespan and resource allocation for both males and females (Koliada *et al*. 2020).

### Implications for multicellular eukaryotes

Variant tRNA-induced mistranslation affects a wide range of physiological processes and exerts both positive and negative effects on flies. Mistranslation is most deleterious during periods of intense growth and translational activity, including embryogenesis and pupation (Mitchell *et al*. 1977; Mitchell and Petersen 1981; Trumbly and Jarry 1983; Qin *et al*. 2007). However, having reached the adult stage, mistranslating flies demonstrated increased lifespan and climbing performance. Previous studies examining mistranslation in complex eukaryotes such as mice, flies, and zebrafish identified developmental defects, organ pathologies, and neurodegeneration, but did not report beneficial effects (Lee *et al*. 2006; Lu *et al*. 2014; Reverendo *et al*. 2014; Liu *et al*. 2014). In contrast to our results, Lu *et al*. (2014) found that male flies constitutively expressing an editing-defective PheRS experience reduced lifespan and climbing ability. These differences may reflect differences in amino acid substitution or level of mistranslation. Some stress conditions, such as heat or cold shock, provide long-term resistance to future stresses after exposure (Hercus *et al*. 2003; Rattan 2005; Le Bourg 2007). Low levels of mistranslation, as we induced with the tRNA variants, may provide similar hormetic effects with physiological benefits. This idea is supported by the V➔S variant increasing lifespan the most despite causing the lowest amount of mistranslation.

We acknowledge that there may be survivorship bias occurring with our mistranslating lines as the adult flies used for longevity and climbing assays necessarily escaped death during development. However, when comparing lifespan of the top 25% longest-lived females from mistranslating or control lines, both T➔S and V➔S females still live significantly longer than corresponding control females (Supplemental Figure S4). This observation suggests that the increase in mistranslating female lifespan is not primarily caused by death of unfit flies prior to adulthood. In the future, development of an inducible system to activate mistranslation specifically during adulthood could explore this potential bias.

We recognize that fruit flies and mammals cope with proteotoxic stress differently, but given that mistranslation is associated with disease (Lant *et al*. 2019) and neurons are expected to be especially vulnerable to translation errors (Drummond and Wilke 2008), our result that two types of mistranslation improve fly locomotion and do not cause neurodegeneration as observed in climbing and rough-eye assays is surprising. Supporting our results, expressing various mistranslating tRNAs in mouse of human neuroblastoma cell lines did not significantly increase cell death (Lant *et al*. 2021; Hasan *et al*. 2023; Davey-Young *et al*. 2024). These results indicate that some forms of tRNA-induced mistranslation are well-tolerated by neuronal cells and may confer protective effects.

## ACKNOWLEDGEMENTS

We would like to thank Dr. Patrick O’Donoghue, Dr. Robert Cumming, Dr. Jiqiang Ling, Dr. Greg Gloor, Dr. Yolanda Morbey and Ecaterina Cozma for their feedback and guidance on the manuscript. This work was supported by Natural Sciences and Engineering Research Council of Canada (NSERC) grants to CJB [RGPIN-2020-07046] and AJM [RGPIN-2020-06464], NIH grants to JV [R01AG056359, R56AG049494 and R35GM119536], as well as a University of Western Ontario Medical & Health Science Research Board seed grant to AJM. JRI was supported by an NSERC Postgraduate Scholarship (Doctoral).

